# Darting across space and time: Parametric modulators of sex-biased conditioned fear responses

**DOI:** 10.1101/2021.06.30.450556

**Authors:** Julia R. Mitchell, Sean G. Trettel, Anna J. Li, Sierra Wasielewski, Kylie A. Huckleberry, Michaela Fanikos, Emily Golden, Mikaela A. Laine, Rebecca M. Shansky

## Abstract

Pavlovian fear conditioning is a widely used behavioral paradigm for studying associative learning in rodents. Despite early recognition that subjects may engage in a variety of both conditioned and unconditioned responses, the last several decades have seen the field narrow its focus to measure freezing as the sole indicator of conditioned fear. We previously reported that female rats were more likely than males to engage in darting, an escape-like conditioned response that is associated with heightened shock reactivity. To determine how experimental parameters contribute to the frequency of darting in both males and females, we manipulated factors such as chamber size, shock intensity, and number of trials. To better capture fear-related behavioral repertoires in our animals, we developed ScaredyRat, an open-source custom Python tool that analyzes Noldus Ethovision-generated raw data files to identify Darters and quantify both conditioned and unconditioned responses. We find that like freezing, conditioned darting occurrences scale with experimental alterations. While most darting occurs in females, we find that with an extended training protocol, darting can emerge in males as well. Collectively, our data suggest that darting reflects a behavioral switch in conditioned responding that is a product of both an individual animal’s sex, shock reactivity, and experimental parameters, underscoring the need for careful consideration of sex as a biological variable in classic learning paradigms.

## Introduction

Pavlovian fear conditioning is an experimental paradigm used to study associative learning in rodents (Fanselow 1984; Maren 2001; Frankland et al. 2004). A neutral conditioned stimulus (CS)— usually an auditory tone—is paired with an aversive unconditioned stimulus (US), usually an electric foot shock. The US evokes an unconditioned response (UR) in the rodent, which learns to associate the CS with the US and ultimately exhibits a conditioned response (CR) when presented with the CS. The most commonly studied CR in Pavlovian fear conditioning is freezing, defined as the total lack of all movement except that required by respiration (Fanselow 1980). For decades, Pavlovian fear conditioning studies have relied solely on the amount of time an animal spends freezing as an indicator of both the level of fear that the animal is experiencing, and of the strength of the CS-US association. This reliance on a single behavior excludes from analysis any other CRs that the animal might engage.

In 2015, our lab identified another CR: darting (Gruene et al. 2015). Darting is characterized by a quick movement across a fear conditioning chamber and occurs in about 40% of female rats and about 10% of males. Using a seven CS-US auditory fear conditioning paradigm, we found that darting emerges during the later tones of the trial, often following earlier freezing responses. This pattern suggests that darting reflects a switch in conditioned responding that is more common in females. Since this initial report, other labs (and our own) have corroborated the finding that conditioned darting occurs primarily in female rats (Pellman et al. 2017; Colom-Lapetina et al. 2019; Greiner et al. 2019; Morena et al. 2021), but the significance of darting as a sex-biased conditioned fear response and the factors that modulate it have not been studied systematically. In contrast, a great deal of work has been done to examine the factors that affect conditioned freezing, including shock intensity (Fanselow 1982), context cues (Bolles and Collier 1976; Fanselow 1980), amount of pre-exposure to the context (Fanselow 1990), length of CS (Fanselow et al. 2019), and number of CS-US pairings (Maren 1998). However, these seminal studies did not consider the sex of the subjects as a potentially data-driving variable.

The goal of the current study was to determine the phenotypic scope and situational modulators of conditioned darting in both males and females. In other words, we asked whether the propensity to dart is pre-determined (i.e. will always occur in the same proportion of males and females), or whether—like freezing—it can be manipulated by altering experimental parameters. We therefore investigated the influence of space, time, and shock intensity on conditioned fear behaviors in both male and female rats. To perform automated, unbiased detection of darting, freezing, and shock reactivity, our lab developed ScaredyRat, an open-access, custom Python tool that analyzes Noldus Ethovision-generated raw data files to identify Darters and quantify behavior throughout a trial. Our findings advance our previous work by illuminating both the experimental conditions under which darting is more or less likely to occur, as well as individual behavioral correlates of darting in both males and females. Overall, this work broadens the field’s understanding of the multiple ways that male and female rodents can show conditioned fear, and underscores the need for more diverse behavioral considerations in classic learning paradigms (Shansky 2018).

### Introducing ScaredyRat

ScaredyRat is an open source, custom Python tool that we developed to assist Ethovision users in evaluating darting, freezing, and shock response behavior in rodent fear conditioning experiments. Graphical User Interfaces (GUIs) for both Windows and Mac users, detailed user manuals, as well as code for Python-proficient users can be found at https://github.com/trettels/ScaredyRat_ver2. ScaredyRat processes raw data files exported from Ethovision and generates spreadsheets for freezing, darting, and velocity metrics, organized according to user-defined Epochs within an experimental session. These Epochs can include, but are not limited to: CS presentation, US presentation, and any pre- or post-Epoch periods of time in which behavior may be of interest. In addition, ScaredyRat generates individual velocity plots (Figure 1) for each subject with color-coded Epochs and bouts of darting and freezing indicated. We note that ScaredyRat as currently designed can only process Ethovision data files. However, our open-source code is available for users who wish to adapt it to be used with other locomotion-tracking programs (e.g. AnyMaze).

**Figure 1.**
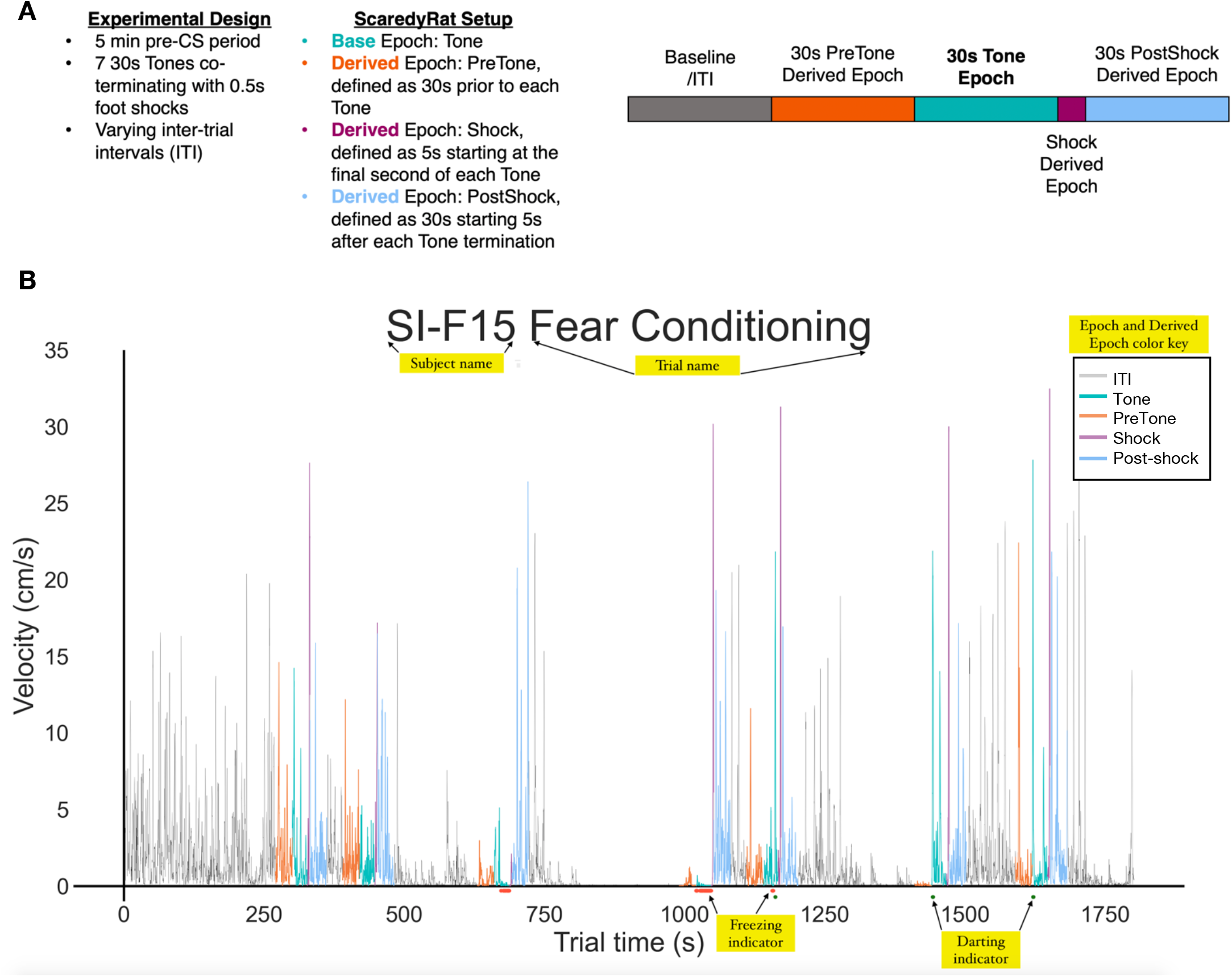
How does ScaredyRat work? A) Example Experimental Design, ScaredyRat setup, and Epoch/Derived Epoch schematic for the plot generated in B). Our experimental design consisted of a 5 min exploration period, followed by 5 30s Tones that co-terminated with a 0.5s footshock, with varying ITIs. ScaredyRat identifies Ethovision-defined Base Epochs (here, the Tone), and then the ScaredyRat user defines Derived Epochs of interest that are time-linked to the Base Epoch (here, PreTone, Shock, and PostShock). B) Representative ScaredyRat-generated plot for a single subject across time in an entire fear conditioning session. Yellow labels indicate notable features—these labels do not appear in actual plots. This animal would be classified as a Darter, because CS (tone)-related darting can be observed in CS 5, 6, and 7. Note earlier periods of freezing on CS 3 and 4. ScaredyRat also generates spreadsheets for Freezing, Darting, and Velocity during each Epoch and Derived Epoch. Behavior during ITIs is not reported.

## Results

We used ScaredyRat to collect freezing, darting, and shock response data in a series of experiments designed to a) identify experimental conditions that may promote or suppress darting, and b) determine the relationship between these three behavioral metrics in both male and female rats.

### 1. Darting requires shock exposure

We began the current investigation by replicating our previous 7 CS-US auditory cued fear conditioning experiment in cohorts of male (n=12) and female (n=12) Sprague Dawley rats (Figure 2A). As in our prior work (Gruene et al. 2015; Colom-Lapetina et al. 2019), only darts that occurred during the CS were used to classify animals as Darters, and we once again found that females comprised the majority of Darters (Figure 2C). Also similar to these past reports, females exhibited greater shock responses than males (main effect of sex: F_1,22_=8.2, p=0.009; Fig 2B, left), a sex difference that appears to have been driven by Darters (main effect of group: F_2,21_=5.5, p=0.01; adjusted post-hoc Darters vs. non-darter Males p=0.009; Fig 2B, right). Although there were no overall sex differences in CS-elicited freezing during acquisition (Fig 2D, left), a 1-way ANOVA of average freezing in Males, Darters, and Non-darter Females produced a significant main effect of group (F_2,21_=3.5, p<0.05) with Darters freezing significantly less than non-darting Males (adjusted post-hoc p=0.03). No sex differences or darting effects were observed on Day 2 (Fig 2E).

**Figure 2.**
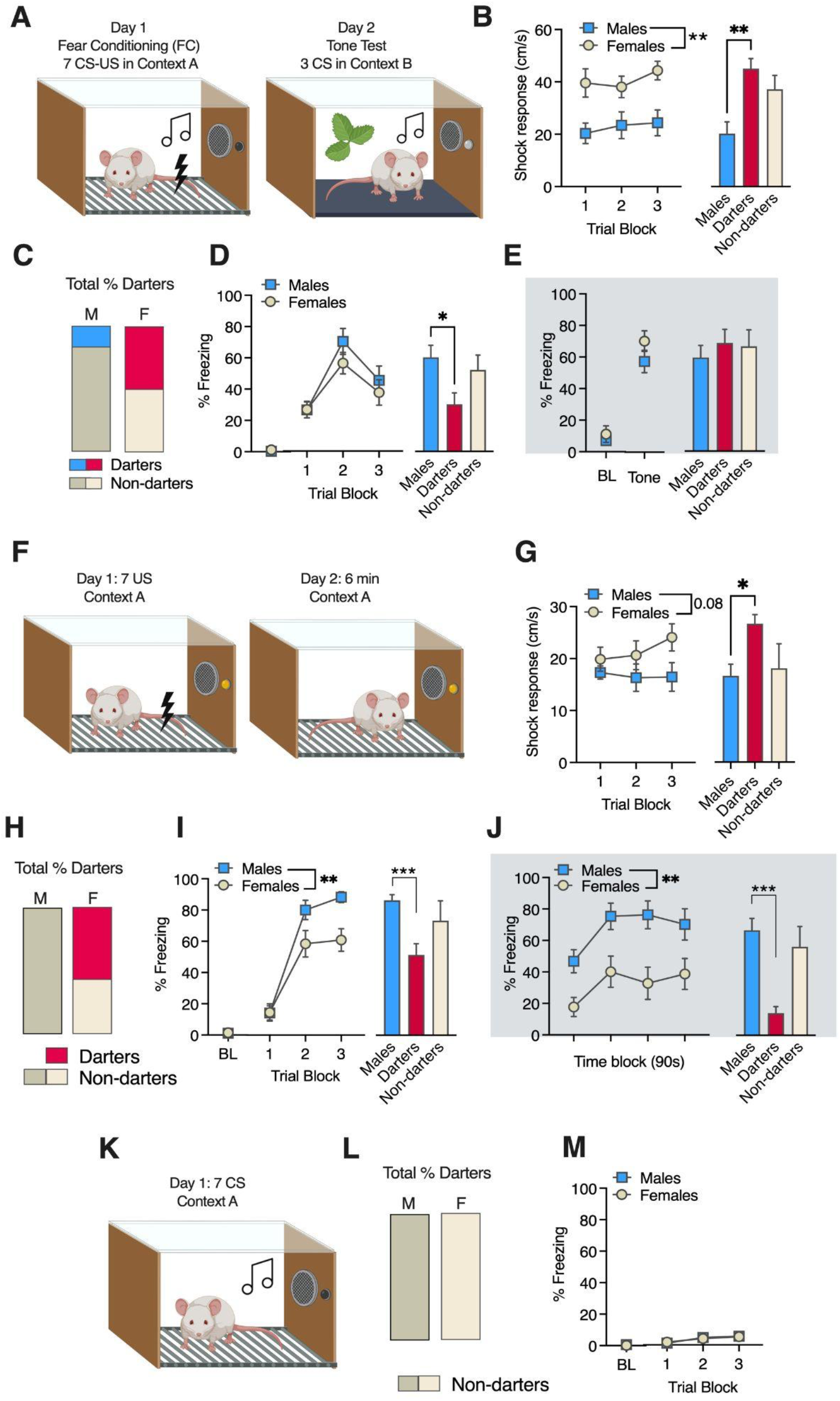
Darting occurs in both cued and context conditioning experiments. A) Experimental design for cued conditioning. B) Shock response in Males and Females across trial blocks (left) and on average (right) with Darters (fuchsia) shown separately from non-darting Males (blue) and females (white). C) Proportions of Darters in each sex (n=12 each sex) D) No sex differences in freezing during conditioning were observed (left), but Darters froze less than non-darting Males. E) No group differences in freezing were observed on Day 2. F) Experimental design for context conditioning. G) Overall sex differences in shock response across trials did not reach significance (left), but analysis by Darting status showed that Darters exhibited higher shock responses than non-darting Males. H) Proportions of Darters in each sex (n=13 males, n=14 females) I) Sex differences in freezing during conditioning (left) were driven by Darters (right). J) Sex differences in freezing during the memory test were driven by Darters, who froze less than Males. *p<0.05, ** p<0.01 *** p<0.001. Comparisons indicated by brackets. K) Experimental design for CS-only conditioning. L) No Darters were observed. M) CS-elicited freezing was minimal in both sexes. Fear conditioning Trial Blocks (Panels D, H, M)) represented as averages of Trials 1-2, 3-4, 5-7. BL: baseline freezing, defined as the first 2 minutes of the session. Panels A, F, K created with BioRender.

To investigate whether darting could be observed in the absence of a discrete CS, we next performed a 2-day context conditioning experiment, in which adult Male (n=12) and Female (n=14) Sprague Dawley rats were exposed to seven footshocks with varying inter-trial intervals, followed by a 6-minute same context exposure 24 hours later (Figure 2F). Darting for each trial was defined as any instance of a discrete movement that a) reached the velocity threshold for darting (20 cm/s) and b) occurred outside the “Shock” and 30s “PostShock” Derived Epochs (See Fig 1 for Epoch illustration), as we view activity during these epochs as direct responses to the shock (i.e. unconditioned responses), rather than conditioned responses. Animals that exhibited at least one dart as just defined were classified as Darters. With these criteria, 57% of Females were classified as Darters but no Males were (Figure 2H).

We next measured the shock response in all animals, defined as the maximum velocity with which the animal moves at the time of shock delivery (see <u>Behavioral Definitions</u> in **Materials & Methods**). Although Females appeared to increase their shock response as trials progressed (Figure 2G, left), effects of sex on shock response neared but did not reach statistical significance (F_1,24_=3.2, p=0.08). However, when Females were separated into Darters and Non-darters, Darters exhibited a significantly higher average shock response than Males (Fig 2G, right: 1-way ANOVA main effect of group: F_2,23_=3.9, p=0.03; post-hoc Darters vs. Males: p=0.02). Freezing levels for Day 1 were sampled by assessing % time freezing during the 30s prior to each shock presentation, to match the timing of freezing measurements taken in our standard cued conditioning experiments. A 2-way ANOVA of freezing behavior during fear conditioning (Figure 2I, left) revealed a significant time x sex interaction (F_2,48_=5.6, p=0.006), as well as significant main effects of time (F_2,44_=117.2, p<0.0001) and sex (F_1,24_=4.7, p=0.039), with Females freezing significantly less than Males. This sex effect appears to be driven by Darters (Figure 2I, right), as a one-way ANOVA of average freezing in Males, Darters, and Non-darters during Trial Blocks 2-3 revealed a significant group effect (F_2,23_=7.2, p=0.004). An adjusted Dunnett’s post-hoc test resulted in a significant difference between Males and Darters (p=0.002) but not Males and Non-darters (p=0.36).

On Day 2, no darting was observed in either sex. Freezing behavior was assessed across the entire 6-minute session in 90s Time Blocks (Figure 2J, left), and we again found significant main effects of time (F_3,63_=11.7, p<0.0001) and sex (F_1,24_=9.5, p=0.005). To assess the contribution of Darters to this effect, we performed a one-way ANOVA of average freezing across the session (Figure 2F, right) and found a significant group effect (F_2,23_=11.8, p=0.0003). Adjusted Dunnett’s post-hocs once again resulted in a significant difference between Males and Darters (p=0.0002) but not Males and Non-darters (p=0.61). We therefore demonstrate here that the engagement of a single darting response during context fear conditioning is sufficient to identify animals that exhibit increased shock responding during context conditioning and reduced conditioned freezing across multiple days. Putative sex differences in context fear conditioning experiments may therefore be driven by a subset of females that engage in darting behavior during training.

Finally, we asked whether darting could be observed in the absence of a US. Male (n=13) and female (n=10) rats were exposed to 7 CS presentations in an identical chamber and with identical ITIs to the first experiment (Fig 2K). No animals of either sex exhibited darting during CS presentation (Fig 2L). As expected, CS-elicited freezing was extremely low, and no sex differences were observed (Fig 2M).

Together, these data demonstrate that while exposure to a footshock US is necessary to observe conditioned darting, a discrete CS is not. To examine the possibility that darting is simply a random response to the shock and not a specific conditioned response, we generated trial-by-trial representations of darting for all three experiments (Figure 3). We examined the prevalence of darting during the Pre-tone (Exp. 1, Fig 3A-B) or pre-shock epochs (Exp. 2, Fig 3C-D) that preceded trials in which respective CS or ITI periods had high levels of darting. A chi-square test comparing Pre-tone vs. Tone epochs in trials 5-7 of Exp. 1 revealed a significant difference in females (chi-sq=5.3, p=0.02), showing that darting is not reflective of a general increase in locomotor activity. Similarly, ITI darting was significantly more prevalent than in the pre-shock epoch for trials 3-6 in Exp. 2 (chi-sq=8.2, p=0.004). Importantly, darting does not emerge spontaneously in animals that are exposed to the auditory CS without footshock, suggesting that darting is not simply reflective of hyperactivity or startle to the tone.

**Figure 3.**
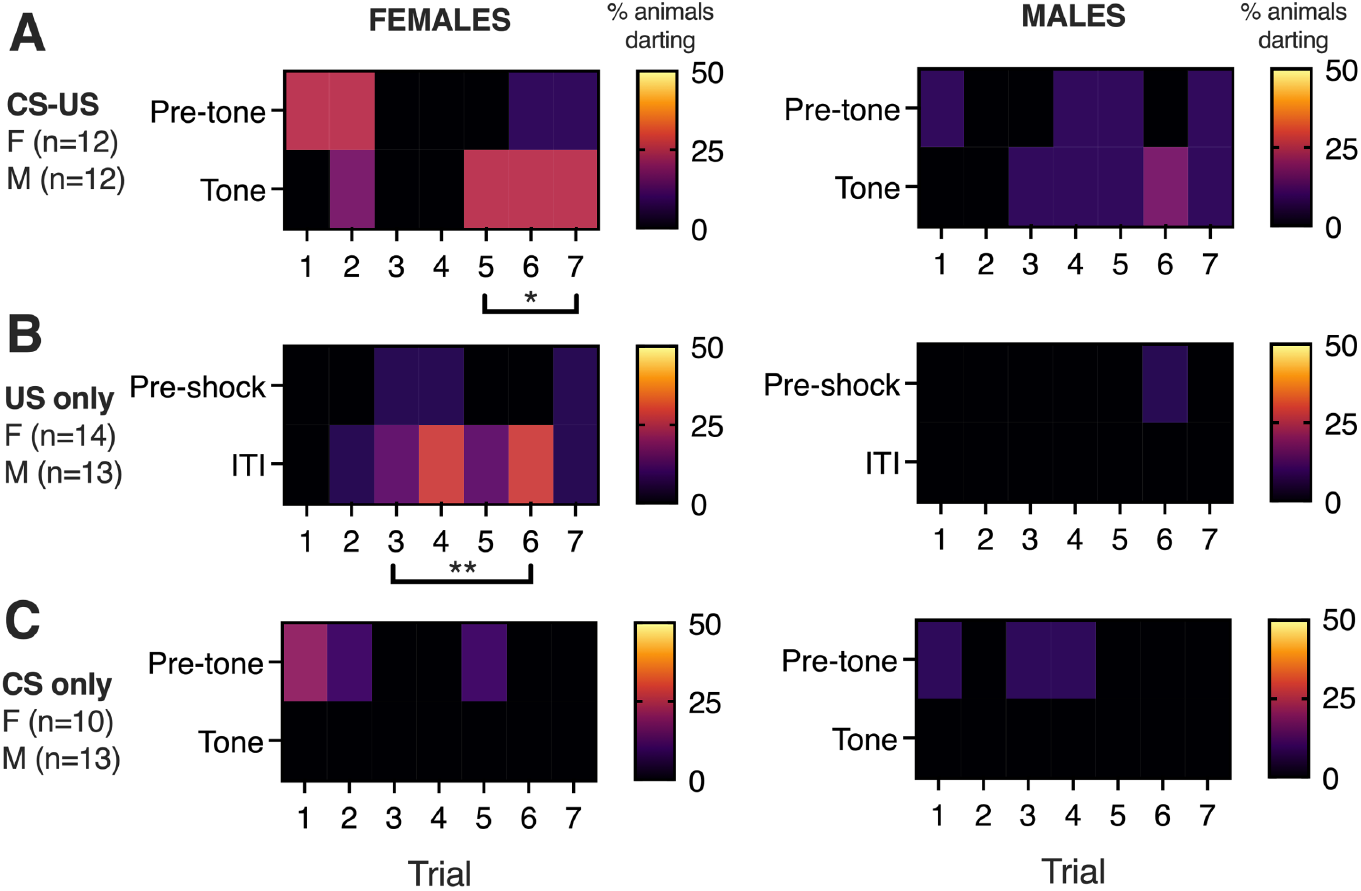
Darting prevalence in Females and Males across trials. Heatmaps for Females (left) and Males (right) show the % of animals that exhibited darting on each trial of the three experiments described above. A) In Exp 1 (cued conditioning), conditioned darting emerged during trials 5-7. We did not observe a comparable increase in locomotor activity during Pre-tone epochs, supporting the idea that darting is a conditioned response. B) In Exp 2 (context conditioning), the emergence of darting was primarily observed during ITIs but not preshock epochs. C) In Exp 3 (CS only), darting in response to the tone was not observed. *p<0.05 **p<0.01 chisq comparing darting in Tone vs Pretone or ITI vs Pre-shock epochs for the indicated trials.

### 2. Darting as a function of chamber size

Rodents can alter conditioned freezing behavior based on the size of the chamber (Bolles and Collier, 1976). Whether darting is also susceptible to such spatial manipulations is not known, nor have potential sex differences in the propensity to adapt conditioned responding as a function of space been investigated. To address these questions, we performed a two-day experiment in which adult Male (n=20) and Female (n=30) Sprague Dawley rats underwent a 7 CS-US cued fear conditioning session in standard 20cm x 20cm chambers (Figure 4A). Twenty-four hours later, they were placed in a 1m^2^ Open Field arena and exposed to seven CS presentations. In half of each cohort, animals’ movement was restricted by an interior arena with 20cm x 20cm dimensions (Small Field, SF), while the other half were given free range of the Open Field arena (OF).

**Figure 4.**
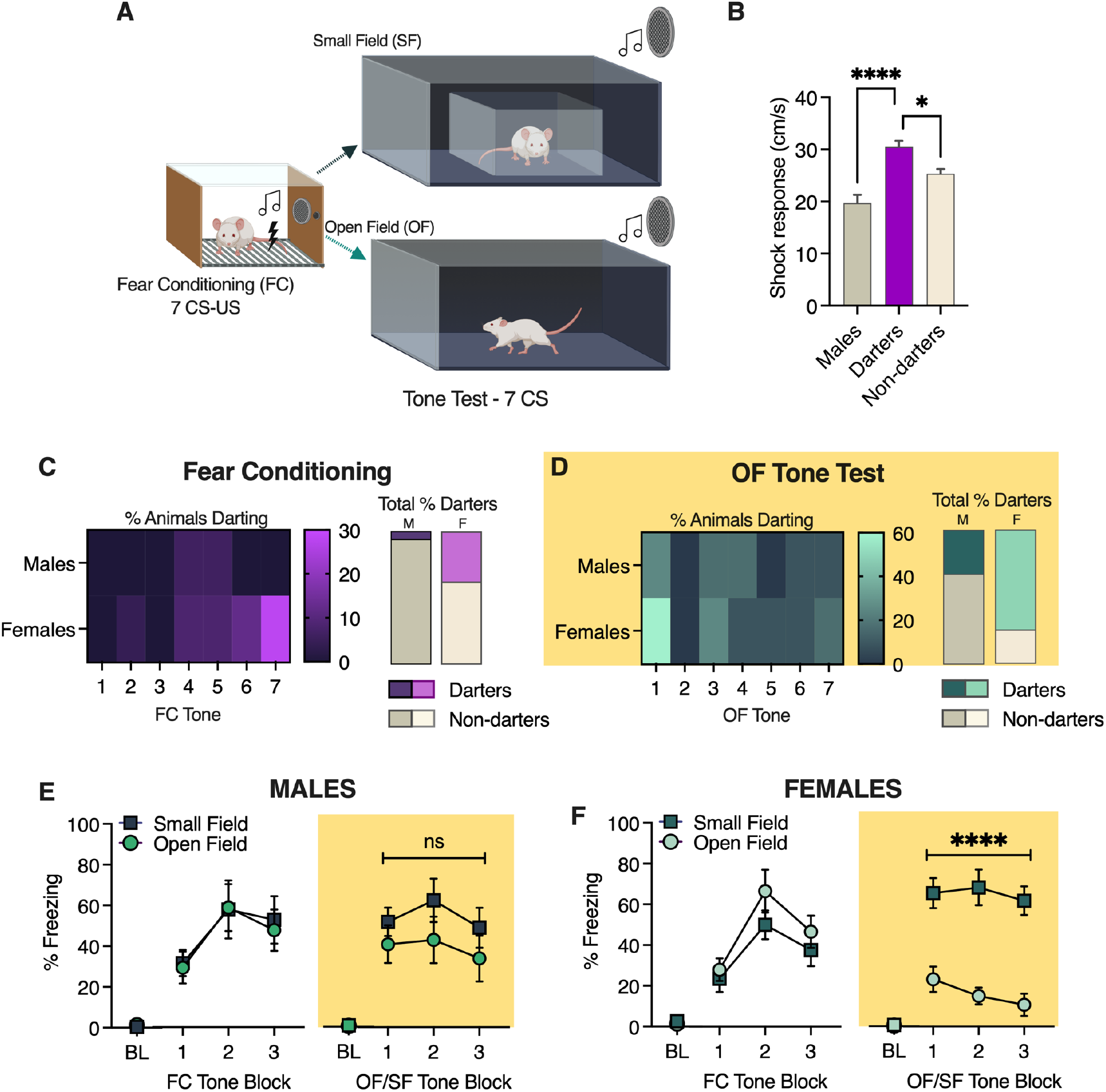
Effects of chamber size on conditioned darting and freezing. A) Experimental design. B) Group differences in shock response on Day 1 for both future SF and OF cohorts C) Darting prevalence during fear conditioning tones. D) Darting prevalence during the tone test in the Open Field. C) Conditioned freezing in Males during fear conditioning (standard chamber) and on Day 2 (either SF or OF). D) Conditioned freezing in Females during fear conditioning (standard chamber) and on Day 2 (either SF or OF).All Tone Blocks (Panels E-F represented as averages of Tones 1-2, 3-4, 5-7. BL: baseline freezing, defined as the first 2 minutes of the session, prior to the first CS. Panel A created with BioRender. ****p<0.01 *p<0.05, comparisons as noted in B), main effect of Field in F).

We first evaluated shock response in all animals on Day 1 and found a significant main effect of group (F_2,46_=13.98, p<0.0001). Dunnett’s multiple comparisons tests revealed that shock response in Darters was significantly higher than both Males (p<0.0001) and Non-darting females (p=0.03; Figure 4B).

Darting across trials and proportion of male and female Darters during Fear conditioning and the Tone test are shown in Figure 4C and D, respectively. As we have found previously, darting during fear conditioning emerges during later trials, and was more prevalent in females compared to males (38% vs. 6%; chi-sq=6.6, p=0.01). On Day 2, we did not observe any conditioned darting in SF Males or Females (data not depicted), consistent with previous observations that in our standard size chambers, darting on Day 2 is extremely rare (Gruene et al. 2015). In contrast, we observed conditioned darting in OF Males and Females during early tones (Figure 4D, left), and once again, darting was more prevalent in females compared to males (75% vs. 33%, chi-sq=4.9, p=0.03; Figure 4D, right).

Finally, we evaluated conditioned freezing on both days as a factor of SF/OF assignment within each sex (Figure 4E). In Males, no effect of group was observed on either day (Day 1: F_1,14_=0.02, p=0.87; Day 2: F_1,14_=1.3, p=0.27). In contrast, while the two Female groups’ freezing did not differ on Day 1 (F_1,22_=1.8, p=0.27), Females exhibited a robust difference in freezing on Day 2 (Figure 4F; F_1,22_=36.6, p<0.0001), with OF Females exhibiting less freezing than their SF counterparts.

The results here demonstrate that while both Males and Females are more likely to exhibit conditioned darting when the space available to them increases, the female-biased nature of darting persists. Furthermore, our freezing data indicate that Females are much more likely than Males to adapt their conditioned fear behavior according to their environment, an important consideration for future fear conditioning studies in which spatial parameters are manipulated.

### 3. Darting as a function of fear conditioning trial number

We reliably observe that darting appears to emerge as CS-US trials progress—often after brief bouts of conditioned freezing—suggesting a switch in conditioned response strategy (see Figure 1 for example). If this is the case, then the animals that engage in darting during a 7 CS-US session may be “early adopters” — in other words, the first animals to switch—and we may thus observe darting in more animals, including Males, if CS-US presentations continue. To answer this question empirically, we performed a 2-day overtraining fear conditioning experiment (20 CS-US presentations, followed by 2 CS presentations in a new context on Day 2) in adult Male (n=19) and Female (n=20) Sprague Dawley rats (Figure 5A).

**Figure 5.**
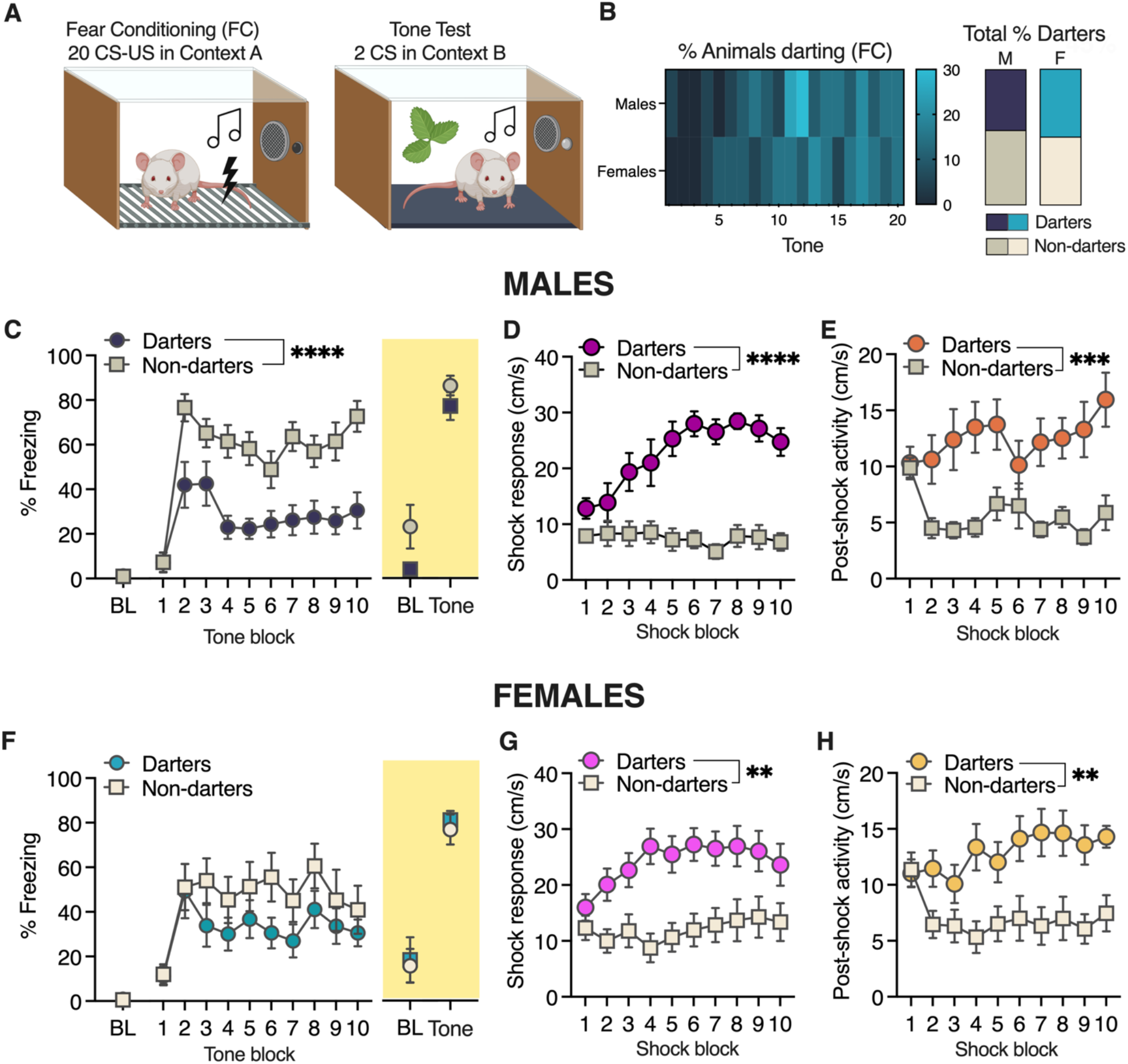
Conditioned darting and freezing in an overtraining paradigm. A) Experimental design. B) Proportion of animals darting during each CS (left) and overall for the entire session C) Conditioned freezing in Male Darters and Nondarters during fear conditioning and on Day 2. D) Shock responses diverged in Male Darters and Non-darters as trials progressed. E) After the first two trials, PostShock activity rapidly diverged in Male Darters and Non-darters. F) Conditioned freezing in Females during fear conditioning and on Day 2. G) Shock responses diverged in Female Darters and Non-darters as trials progressed. H) After the first two trials, PostShock activity rapidly diverged in Female Darters and Non-darters. All statistical significance noted represents within sex Darters vs. Non-darters. ****p<0.0001, **p<0.01. Panel A created with BioRender. BL: baseline freezing, defined as the first 2 minutes of the session, pre-CS. All Tone and Shock blocks represented as averages of every two trials.

When we evaluated the prevalence of darting across fear conditioning trials (Figure 5B, left), we observed that as in previous studies, darting in Females emerged around CS 5-7. In Males, we observed an increase in darting shortly after that, with nearly 30% of Males darting during tones 11-12. Overall, 45% of Males and 50% of Females were classified as Darters (Figure 5B, right), suggesting that sex differences in darting prevalence can be eliminated with extended protocols.

The near 50/50 split in both sexes allowed us to conduct within-sex comparisons of other behaviors in Darters vs. Non-darters. In Males, Darters exhibited reduced freezing during fear conditioning but not the tone test on Day 2 (Figure 5C). A 2-way ANOVA for freezing on Day 1 revealed a significant darting x trial interaction (F_9,153_=2.1, p=0.04) and a significant main effect of darting (F_1,17_=26.1, p<0.0001). Analysis of shock responding (Figure 5D) also revealed a significant darting x trial interaction (F_9,153_=6.9, p<0.0001) and a significant main effect of darting (F_1,17_=42.0, p<0.0001), driven by an increase in shock response velocity in darters. Similarly, evaluation of PostShock activity (Figure 5E), defined as the maximum velocity reached in the 30s PostShock Derived Epoch (see Figure 1), revealed a significant darting x trial interaction (F_9,153_=2.6, p=0.008) and a significant main effect of darting (F_1,17_=16.3, p=0.0009).

Surprisingly, Female Darters and Non-darters did not significantly differ in conditioned freezing during fear conditioning or the tone test (Figure 5F; 2-way ANOVA effect of darting: F_1,18_=1.87, p=0.19). However, analysis of shock responding (Figure 5G) revealed a significant darting x trial interaction (F_9,162_=2.6, p=0.007) and a significant main effect of darting (F_1,18_=10.8, p=0.004). As in Males, this effect was driven by a progressive increase in Female Darters’ shock response velocity. We observed a similar pattern in analysis of PostShock activity (Figure 5H), finding a significant darting x trial interaction (F_9,162_=2.3, p=0.02) and a significant main effect of darting (F_1,18_=12.0, p=0.003).

Interestingly, the significant effect of darting in both Males and Female PostShock responding appears to be primarily driven by a rapid drop in PostShock activity exhibited by Non-darters after the first two shocks, while Darters maintain or increase their activity levels.

We conclude from these analyses that with an extended or “overtraining” fear conditioning session, similar proportions of Males and Females will engage in conditioned darting. These findings support the idea that darting reflects a switch in conditioned fear behavior. Moreover, darting in both sexes is correlated with an increase in shock response and PostShock locomotor activity, suggesting that individual differences in shock sensitivity may be an important predictor of an animal’s propensity to dart.

### 4. Darting as a function of shock intensity

Our previous and current results suggest that darting may arise in animals that exhibit an amplified and protracted behavioral response to the shock. To investigate the role that shock intensity might play in modulating the prevalence of darting, we performed a 2-day auditory cued fear conditioning experiment in which animals (total Males: n=24; total Females: n=23) were exposed to seven presentations of our standard 30-s CS, which co-terminated with either a 0.3mA or 1.0mA foot shock (Figure 6A). A two-CS tone test was performed in a new context the following day.

**Figure 6.**
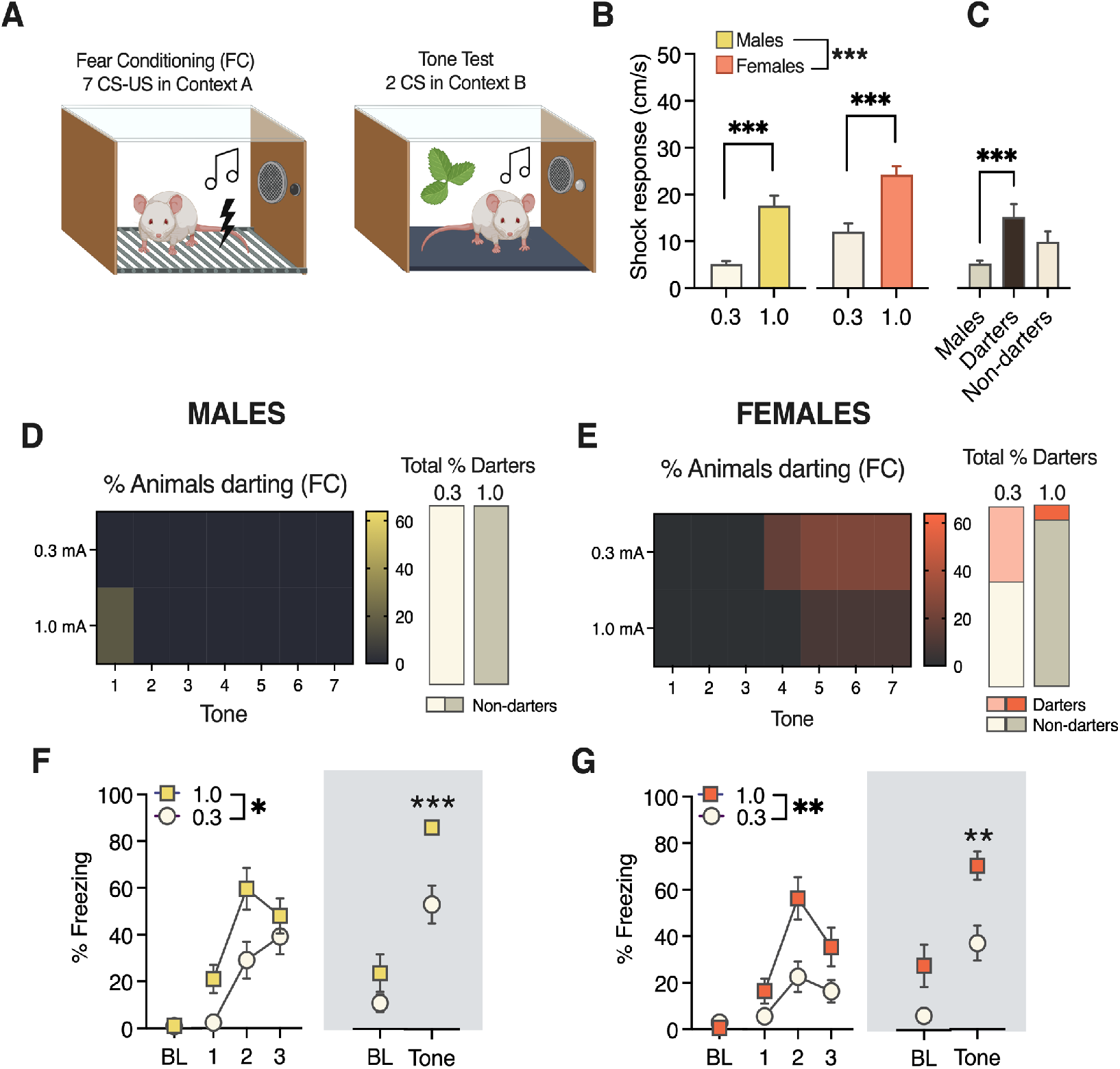
Conditioned darting and freezing as a function of shock intensity. A) Experimental design. B) Average shock response in Males and Females for 0.3mA and 1.0mA experiments. C) Average shock response for 0.3mA separated by Darters. D) Proportion of Males that darted during each CS (left) and overall for the entire fear conditioning session (right). E) Proportion of Females that darted during each CS (left) and overall for the entire fear conditioning session (right). F) Freezing in Males during fear conditioning (left) and a 2-CS test in a new context 24h later. G) Freezing in Females during fear conditioning (left) and a 2-CS test in a new context 24h later. Unless specifically noted (B), all statistical significance symbols represent within sex differences between 0.3mA and 1.0mA groups. *p<0.05, **p<0.01, ***p<0.001. Panel A created with BioRender. BL: baseline freezing, defined as the first 2 minutes of the session, pre-CS 1. All fear conditioning Tone Blocks (Panels F-G [left]) represented as averages of Tones 1-2, 3-4, 5-7. CS-elicited freezing on Day 2 is represented as the average of 2 CSs. All group n’s=12.

Analysis of average shock response in all animals revealed main effects of both sex and shock intensity (sex: F_1,44_=15.98, p=0.0002; shock: F_1,44_=15.98, p<0.0001; Figure 6B). Sidak’s adjusted post-hoc tests confirmed that both Males and Females exhibited greater shock responses to the 1.0mA shock compared to the 0.3mA shock (p<0.0001 both comparisons) and that Females exhibited more robust shock responses than males at both shock intensities (p<0.05 both comparisons). We also performed a 1-way ANOVA of shock response in the 0.3mA groups separated by darting, and found as previously, Darters’ shock response was significantly higher than Males’ (Figure 6C; F_2,21_=9.3, p=0.001; Dunnett’s post-hoc p=0.0008). Non-darters did not differ from Males (p=0.07), suggesting that the main sex difference was driven at least in part by Darters. We did not perform this analysis in the 1.0mA condition due to the very low number of Darters (Fig 6E).

Darting prevalence in Males and Females across trials and overall can be seen in Figure 6D and 6E, respectively. No Males reached criterion for conditioned darting in either shock condition. In Females, darting emerged during later trials, as previously observed. Five females in the 0.3mA group and one female in the 1.0mA group reached criterion for darting. Although this would suggest that darting is more likely to occur at lower shock intensities, this difference did not reach statistical significance, possibly due to low power (chi-sq=3.2, p=0.08).

Males showed higher levels of CS-elicited freezing when the US was 1.0mA compared to 0.3mA (Figure 6F, left; main effect of intensity: F_1,22_=7.4, p=0.01). This discrimination was also evident in CS-elicited freezing on Day 2 (Figure 5F, right; Mann Whitney U=16; p=0.0007). In Females (Figure 6G), animals in the 1.0mA group also exhibited higher levels of CS-elicited freezing compared to 0.3mA on both days (Day 1, effect of shock: F_1,22_=8.4, p=0.008; Day 2, unpaired t-test = 3.4, p=0.002).

Together, these data tell an intriguing and somewhat paradoxical story about the interactions between shock intensity, shock response, conditioned freezing, and the propensity to engage darting behavior. We found that shock response is a reliable measure to demonstrate discrimination between shocks of different intensities in both sexes, and that shock response in Darters is higher than non-darting Males. However, we also found that CS-elicited darting may be more likely to be observed when shock intensities are *lower*. We discuss the potential implications of these data below and look forward to investigating the mechanistic basis of these surprising findings in future work.

## Discussion

The purpose of these experiments was to identify situational modulators and behavioral correlates of conditioned darting in Male and Female rats during Pavlovian fear conditioning. Manipulations of context, trial number, and shock intensity revealed that, like freezing, darting as a conditioned response is indeed sensitive to these experimental parameters; more space or CS-US trials led to an increase in the number of Darters, while increasing shock intensity decreased the number of Darters. ScaredyRat analysis also demonstrated that Darters of both sexes reliably show higher shock responses and protracted PostShock activity than Non-darters, suggesting that the propensity of an individual animal to engage in darting as a conditioned response is related to the magnitude of its unconditioned responses. Intriguingly, these phenotypic splits in shock reactivity are not evident in the very first trials, suggesting that the initial CS and US experiences may trigger these divergent trajectories. We look forward to more thoroughly investigating the mechanisms that connect pain sensitivity and conditioned responses in the future.

Our findings build on decades of seminal single-sex work that defined how experimental parameters affect conditioned freezing, but we also make significant advances in several notable ways. First, ScaredyRat allowed us to examine not only freezing but also multiple conditioned and unconditioned fear behaviors across different experimental preparations. While freezing is a well-established and dependable conditioned fear response, the field’s current reliance on it as the sole indicator of conditioned fear in rodents has given us an incomplete picture. For decades, any nonfreezing behavior exhibited during classical fear conditioning has been excluded from analyses as not fear-related or not representative of learning. We see here that at least one of those behaviors, darting, is representative of learning and is a conditioned response that is affected by similar situational modulators that affect freezing.

Evidence that darting is a true conditioned response comes from several points: first, we replicated our prior findings (Gruene et al. 2015; Colom-Lapetina et al. 2019) that darting in a cued conditioning experiment follows an upward trajectory as the session progresses (like freezing), occurs only in response to the CS, and does not reflect a general increase in locomotor activity (Figure 3A). In addition, we showed that darting does not occur in response to the CS when the CS does not cooccur with_a US (Figure 3C), demonstrating that it is not an innate response to the CS. We also note that within a CS, darting occurs on average 7-10 seconds after CS onset (Supplemental Figure 1B), suggesting that darting does not reflect a CS-elicited startle response. Further support comes from other labs, who have reported conditioned darting in several rodent strains, suggesting it may be a conserved response. In addition to recent reports of conditioned darting in female Sprague Dawley rats (Morena et al. 2021), female Long Evans rats in a cue-discrimination paradigm (for reward, safety and threat) selectively elicit darting to the threat-paired cue, a behavior that emerges in later trials. Similar to our findings, these animals also show a stronger motor response to the shock itself (Greiner et al. 2019). Furthermore, Long Evans rats (Totty et al. 2021), and both FVBB6 F1 hybrid and C57BL/6J mice (Borkar et al. 2020; Fadok et al. 2017; Hersman et al. 2020) display darting-like active flight behaviors in response to conditioning with a serial compound stimulus, a behavior that is amenable to extinction training. Reports in which darting is not observed (Tryon et al. 2021) may be explained by brevity of conditioning (3 CS-US pairings), and the intensity of the US (1 mA). As we report here, darting typically emerges on or after the 5th tone presentation and occurs less frequently in animals receiving 1 mA shocks compared to milder ones.

Some of the more thought-provoking aspects of our work arise when trying to reconcile the relationship between darting and shock reactivity. In Experiment 3, Darters of both sexes had a higher maximum shock response velocity than non-darters, suggesting that there may be a difference in pain sensitivity between Darters and Non-darters. We might then expect to observe more darting at higher shock intensities, which also elicit higher shock responses. However, Experiment 4 showed that when shock intensity increased, rats were *less* likely to dart, despite clear evidence that the shock was perceived as more intense. Bolles & Fanselow’s perceptual-defensive-recuperative model of pain (Bolles and Fanselow 1980) states that when fearful and painful stimuli are both present, fear will “win out” over pain, leading to an opioid-mediated analgesic effect that allows the animal to respond to the fear-inducing stimulus (e.g., freeze). In this context, one possible explanation of our data is that darting indicates that conditioned analgesia has failed to occur. Supporting this hypothesis is evidence from Experiment 3, in which we found that shock responding in Non-darters stayed relatively flat, while progressively escalating in Darters. Given known sex differences in pain processing and opioid signaling (Loyd et al. 2008; Fullerton et al. 2018; Mogil 2020), we feel that future investigations into the neural circuits that might drive this possibility are likely to be fruitful. In particular, the periaqueductal gray (PAG) is a center for both pain and defensive responding in the rodent (de Oca et al. 1998; Vianna et al. 2001; Vanegas and Schaible 2004) and is an attractive target for mechanistic inquiry.

This work advances and expands upon decades of research on conditioned fear, the vast majority of which has been conducted only in males and only reports freezing behavior. Although landmark papers from the 1970s and 1980s were more likely to include evaluation of both freezing and shock-related activity metrics (e.g. activity bursts or circa-strike behavior; reviewed in (Fanselow 1994), this practice has fallen out of fashion in more recent years, we believe to the detriment of the field of Behavioral Neuroscience. Side-by-side assessment of conditioned fear in both <u>f</u>emales and <u>m</u>ales is still woefully rare (Lebron-Milad and Milad 2012), and we believe that the inclusion of both sexes in conditioned fear inquiries will be critical to capturing the range of conditioned behaviors that may be exhibited, not simply those that are dominant in one sex (Shansky and Woolley 2016; Shansky 2019). Our work provides not only novel insight into the facets of the paradigm in which the sexes do and do not differ but also a novel tool with which to collect data for multiple behavioral outcomes. We hope that as others move to include both sexes in their experimental designs, they will take advantage of ScaredyRat to analyze and identify individual differences in both unconditioned and conditioned fear responses.

## Materials and Methods

### Subjects

All experiments were conducted in young adult (8-10 weeks) Male (n=95) and Female (n=105) Sprague Dawley rats (Charles River), weighing 325-350g and 225-250g, respectively. See **Experimental Design Table** for group n’s. Animals were same sex pair-housed in the Nightingale Hall Animal Facility at Northeastern University in a 12:12 light:dark cycle with access to food and water *ad libitum*, and were allowed to acclimate to the facilities undisturbed for at least one week prior to testing. Male and female animals were housed in the same room. Animals were briefly acclimated to handling for two days prior to testing. Testing was conducted in the light phase between 10am and 4pm in behavior rooms located within 50 ft of the vivarium, so transport was minimal. All experimenters were female. The estrous cycle was not monitored, as we have previously reported that the propensity to dart is not related to the cycle phase (Gruene et. al, 2015). We note therefore that concerns that daily swabbing could induce darting behavior are irrelevant here. All procedures were conducted in accordance with the National Institutes of Health Guide for the Care and Use of Laboratory Animals and were approved by the Northeastern University Institutional Animal Care And Use Committee.

### Behavioral Apparatus and Fixed Experimental Parameters

Fear conditioning training (FC) and recall tone testing (except in Experiment 2) were conducted as described in Gruene et al., 2015, in one of eight identical 20cm^2^ chambers constructed of aluminum and Plexiglass walls (Rat Test Cage; Coulbourn Instruments, Allentown Pennsylvania) with metal stainless steel rod flooring attached to a shock generator (Model H13-15; Coulbourn instruments). Each chamber was enclosed within a sound-isolation cubicle (model H10-24A; Coulbourn Instruments). An overhead, infrared digital camera allowed videotaping (30 frames per second) during behavioral procedures. Chamber grid floors, trays, walls, and ceilings were thoroughly cleaned with water and 70% ethanol and dried between sessions. Chambers were used for both males and females, but test sessions were restricted to a single sex.

For context A, chambers were lit with a single house light. Context B testing was done in the same chambers, but the house light was off and a cue light was illuminated. In addition, plexiglass panels covered the metal grid floor, and a light peppermint scent (Dr. Bronner’s) was applied to a removable tray under the chamber floor. Across experiments, animals were allowed 5 minutes to explore the chamber before CS or US presentation commenced. Mean intertrial interval for all experiments was 3 minutes with a range of 1.5-5 minutes. In all cued fear conditioning experiments, the CS was a 30s, 4 kHz, 80 dB SPL sine wave tone, which co-terminated with a 0.5s footshock. Number of CS presentations and footshock intensity varied for each individual experiment and are noted below in the Experimental Design Table.

### Open Field Arena

For Experiment 2, Day 2 testing occurred in a 1m^2^ wooden Open Field Arena with 30cm height walls, painted with matte black spray paint. For the Small Field condition, a 30cm^2^ wooden box with open top and bottom (also painted with matte black spray paint) was placed inside the Open Field (see Figure 3A). The CS was generated with Garage Band software (Apple) and played at an ITI mimicking those in the standard chambers. Behavior was recorded with a Microsoft Kinect camera.

### Behavior tracking and data processing

We used Ethovision software (Noldus; Leesburg, VA) to generate raw velocity data sheets from all video files at a sample rate of 15 frames per second (i.e. every other frame). These files were then fed to ScaredyRat, which extracted freezing, darting, and velocity data for each animal during defined Epochs and Derived Epochs as conveyed in Results.

### ScaredyRat settings and definitions

- The Time Bin Duration was set at 1s.
- The threshold for **freezing** for all experiments was set at <0.1 cm/s (average speed within a Time Bin).
- The threshold for **darting** for all experiments was set at >20 cm/s.
- **Baseline** (BL) measures reflect the first two minutes of all sessions. No CS presentations occurred during this time.
- **Shock response** is defined as the maximum velocity reached within a 5 second Derived Epoch that begins with the shock presentation.
- **PostShock activity** is defined as the maximum velocity reached within a 30 second Derived Epoch that begins immediately after each Shock Epoch.

### Behavioral definitions

**Freezing** was defined as in (Fanselow 1980)—specifically, the cessation of all movement except for breathing. ScaredyRat does not count freezing bouts under 1s.

**Darting** was defined as in (Gruene et al 2015)—specifically, darts are isolated, discrete locomotor events that reach above 20 cm/s and that do not fall within the immediate shock response epoch. This threshold was determined based on initial human observation of darting—i.e., when hand-scored videos were processed by Ethovision, locomotor events that were scored as darting by at least 2 experimenters all were over 20 cm/s. We distinguish darting from shock response as we believe the shock response to be a reflexive indicator of pain and not a conditioned response.

**Darters** are defined as in (Gruene et al 2015)—specifically, any animal that exhibits one or more darts during at least one CS presentation, excluding CS 1-2. We set this inclusive threshold because in our experience, a single dart is sufficient to identify animals that exhibit behaviors that we have observed previously associated with darting, including enhanced shock response. Answers to common questions about Darters can be found in Figure S1. As we have previously reported, Darters do not differ in weight compared to same-sex Non-darters (unpaired t-tests, Females p=0.72; Males p=0.11; Figure S1A). Examination of dart timing within a CS (in other words, how soon after the CS onset does darting occur?) is shown in S1B (data from Overtraining experiment). The average number of trials during which Darters exhibited darting in each experiment is shown in Figure S1C.

**Supplemental figure 1.**
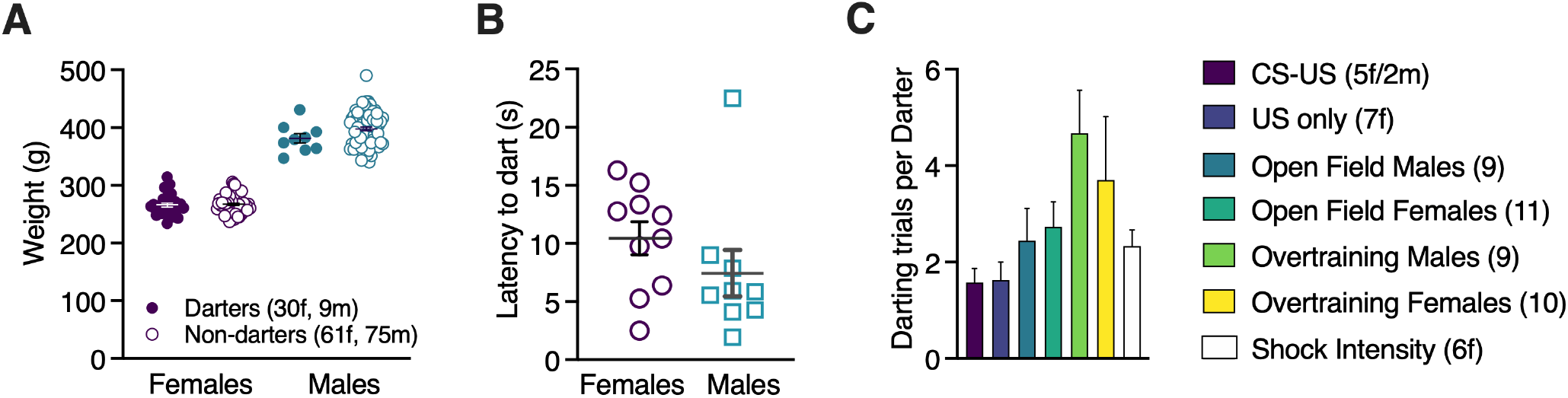
Information about Darters. A) The weights of Female and Male Darters did not differ from Non-darters at the time of behavioral testing. B) Latency to dart after CS onset for male and female Darters in the overtraining experiment, showing that darting is not associated with either immediate CS onset or imminent shock (shock occurs at 30s). C) Average number of trials during which Darters darted for each experiment in this paper. Note that Open Field data are from Day 2.

#### Experimental Design Table

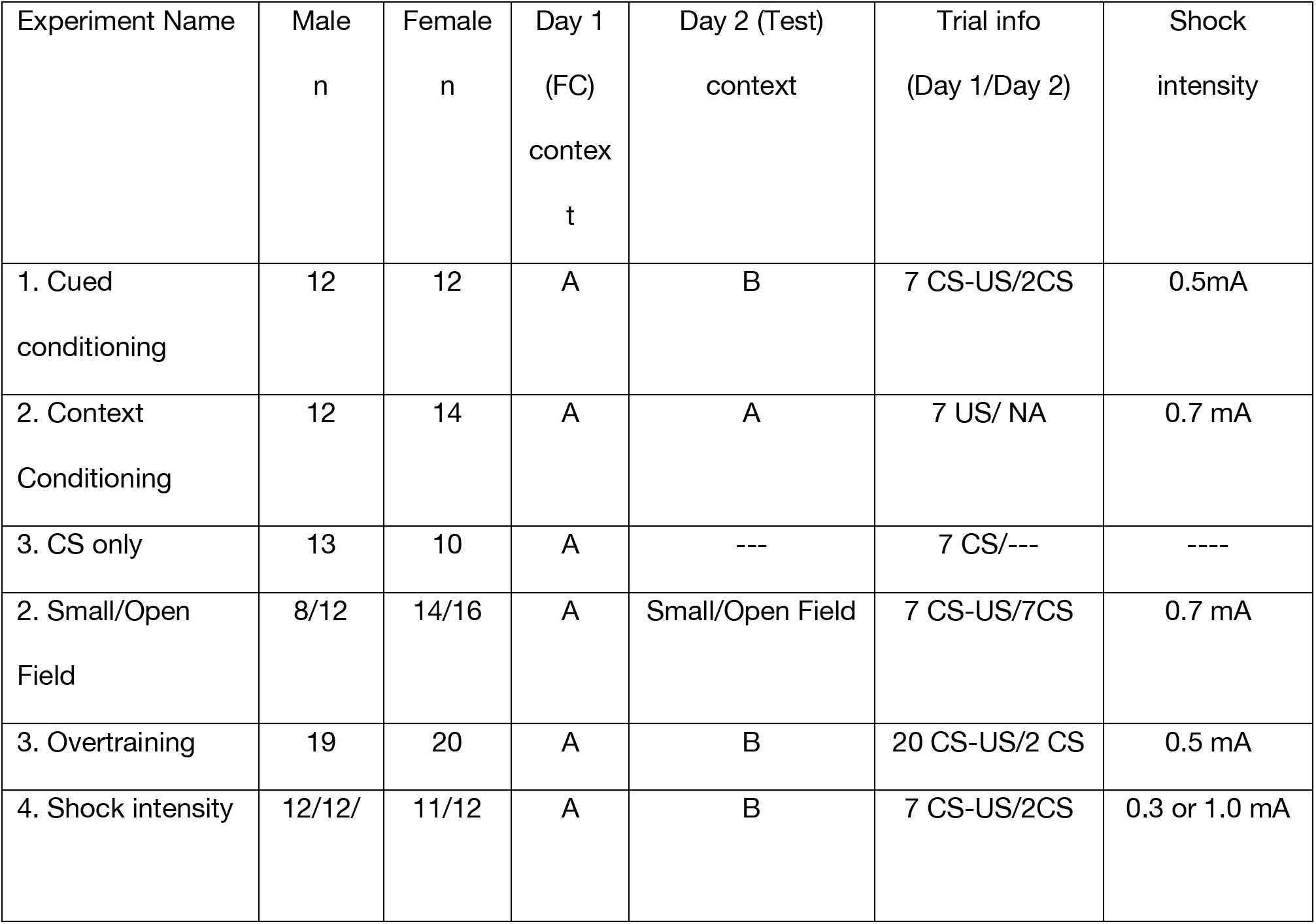

### Statistical analysis

All statistical analyses were done with Graphpad Prism Software. Data sets with multiple time points (e.g., fear conditioning data) were analyzed with 2-way repeated measure ANOVAs, followed by Sidak’s post-hoc tests adjusted for multiple comparisons when appropriate. One-way ANOVAs were followed by Sidak’s or Dunnett’s post-hoc tests when appropriate. Group differences in data sets with a single time point (e.g. Day 2 average freezing) were assessed with either unpaired t-tests or Mann Whitney tests if groups were determined to have unequal variances.

## Acknowledgments

This work was supported by a Fay/Frank Seed grant from the Brain Research Foundation and NIMH R01 MH123803 to RMS. We thank Jesse Schomer and Sandeep “Bob” Datta for providing equipment used in the Open Field experiment, and Vasvi Bhutani for technical assistance with the shock intensity experiment.

## Contributions

JRM, RMS, and SW designed the experiments. AJL, SGT, and KAH developed ScaredyRat. JRM, SW, EG, and MF ran the experiments. JRM and RMS analyzed the data. JRM, MAL, and RMS wrote the manuscript.

## References

Bolles RC, Collier AC. 1976. The effect of predictive cues on freezing in rats. Animal Learning & Behavior 4: 6–8. https://link.springer.com/article/10.3758/BF03211975 (Accessed June 22, 2021).

Bolles RC, Fanselow MS. 1980. A perceptual-defensive-recuperative model of fear and pain. Behavioral and Brain Sciences 3: 291–301. /record/1981-25138-001 (Accessed June 22, 2021).

Borkar CD, Dorofeikova M, Le QSE, Vutukuri R, Vo C, Hereford D, Resendez A, Basavanhalli S, Sifnugel N, Fadok JP. 2020. Sex differences in behavioral responses during a conditioned flight paradigm. Behavioural Brain Research 389.

Colom-Lapetina J, Li AJ, Pelegrina-Perez TC, Shansky RM. 2019. Behavioral diversity across classic rodent models is sex-dependent. Frontiers in Behavioral Neuroscience 13.

de Oca BM, DeCola JP, Maren S, Fanselow MS. 1998. Distinct regions of the periaqueductal gray are involved in the acquisition and expression of defensive responses. The Journal of neuroscience: the official journal of the Society for Neuroscience 18: 3426–32. http://www.ncbi.nlm.nih.gov/pubmed/9547249 (Accessed February 20, 2017).

Fadok JP, Krabbe S, Markovic M, Courtin J, Xu C, Massi L, Botta P, Bylund K, Müller C, Kovacevic A, et al. 2017. A competitive inhibitory circuit for selection of active and passive fear responses. Nature 542.

Fanselow MS. 1980. Conditioned and unconditional components of post-shock freezing. The Pavlovian journal of biological science 15: 177–82. http://www.ncbi.nlm.nih.gov/pubmed/7208128 (Accessed May 31, 2015).

Fanselow MS. 1990. Factors Governing One-Trial Contextual Conditioning. Animal Learning & Behavior 18: 264–270. http://gateway.webofknowledge.com/gateway/Gateway.cgi?GWVersion=2&SrcAuth=mekentosj&SrcApp=Papers&DestLinkType=FullRecord&DestApp=WOS&KeyUT=A1990DQ33800006.

Fanselow MS. 1994. Neural organization of the defensive behavior system responsible for fear. Psychonomic Bulletin & Review 1: 429–438.

Fanselow MS. 1982. The postshock activity burst. Animal Learning & Behavior 10: 448–454. https://link.springer.com/article/10.3758/BF03212284 (Accessed June 22, 2021).

Fanselow MS, Hoffman AN, Zhuravka I. 2019. Timing and the transition between modes in the defensive behavior system. Behavioural Processes 166. https://pubmed.ncbi.nlm.nih.gov/31254627/ (Accessed June 22, 2021).

Frankland PW, Josselyn SA, Anagnostaras SG, Kogan JH, Takahashi E, Silva AJ. 2004. Consolidation of CS and US representations in associative fear conditioning. Hippocampus 14: 557–569. http://eutils.ncbi.nlm.nih.gov/entrez/eutils/elink.fcgi?dbfrom=pubmed&id=15301434&retmode=ref&cmd=prlinks.

Fullerton EF, Doyle HH, Murphy AZ. 2018. Impact of sex on pain and opioid analgesia: a review. Current Opinion in Behavioral Sciences 23: 183–190. /pmc/articles/PMC6428208/?report=abstract (Accessed August 12, 2020).

Greiner EM, Müller I, Norris MR, Ng KH, Sangha S. 2019. Sex differences in fear regulation and reward-seeking behaviors in a fear-safety-reward discrimination task. Behavioural Brain Research 368.

Gruene TM, Flick K, Stefano A, Shea SD, Shansky RM. 2015. Sexually divergent expression of active and passive conditioned fear responses in rats. eLife 4: 1–9. http://eutils.ncbi.nlm.nih.gov/entrez/eutils/elink.fcgi?dbfrom=pubmed&id=26568307&retmode=ref&cmd=prlinks.

Hersman S, Allen D, Hashimoto M, Brito S, Anthony TE. 2020. Stimulus salience determines defensive behaviors elicited by aversively conditioned serial compound auditory stimuli. eLife 9.

Lebron-Milad K, Milad MR. 2012. Sex differences, gonadal hormones and the fear extinction network: implications for anxiety disorders. Biology of Mood & Anxiety Disorders 2012 2:1 2: 1–12. https://biolmoodanxietydisord.biomedcentral.com/articles/10.1186/2045-5380-2-3.

Loyd DR, Wang X, Murphy AZ. 2008. Sex differences in μ-opioid receptor expression in the rat midbrain periaqueductal gray are essential for eliciting sex differences in morphine analgesia. Journal of Neuroscience 28: 14007–14017. https://www.jneurosci.org/content/28/52/14007 (Accessed August 12, 2020).

Maren S. 2001. Neurobiology of Pavlovian fear conditioning. Annual review of neuroscience 24: 897–931. http://www.ncbi.nlm.nih.gov/pubmed/11520922 (Accessed February 16, 2015).

Maren S. 1998. Overtraining does not mitigate contextual fear conditioning deficits produced by neurotoxic lesions of the basolateral amygdala. Journal of Neuroscience 18: 3088–3097. https://www.jneurosci.org/content/18/8/3088 (Accessed June 22, 2021).

Mogil JS. 2020. Qualitative sex differences in pain processing: emerging evidence of a biased literature. Nature Reviews Neuroscience 21: 353–365. https://pubmed.ncbi.nlm.nih.gov/32440016/ (Accessed November 14, 2020).

Morena M, Nastase AS, Santori A, Cravatt BF, Shansky RM, Hill MN. 2021. Sex-dependent effects of endocannabinoid modulation of conditioned fear extinction in rats. British Journal of Pharmacology 178.

Pellman BA, Schuessler BP, Tellakat M, Kim JJ. 2017. Sexually Dimorphic Risk Mitigation Strategies in Rats. eNeuro 4.

S. Fanselow M. 1984. What is conditioned fear? Trends in Neurosciences 7: 460–462.

Shansky RM. 2019. Are hormones a “female problem” for animal research? Science 364: 825–826. http://www.ncbi.nlm.nih.gov/pubmed/31147505 (Accessed June 4, 2019).

Shansky RM. 2018. Sex differences in behavioral strategies: avoiding interpretational pitfalls. Current Opinion in Neurobiology 49.

Shansky RM, Woolley CS. 2016. Considering Sex as a Biological Variable Will Be Valuable for Neuroscience Research. Journal of Neuroscience 36: 11817–11822. http://www.jneurosci.org/content/36/47/11817.abstract.

Totty MS, Warren N, Huddleston I, Ramanathan KR, Ressler RL, Oleksiak CR, Maren S. 2021. Behavioral and brain mechanisms mediating conditioned flight behavior in rats. Scientific Reports 11.

Tryon SC, Sakamoto IM, Kellis DM, Kaigler KF, Wilson MA. 2021. Individual Differences in Conditioned Fear and Extinction in Female Rats. Frontiers in Behavioral Neuroscience 15.

Vanegas H, Schaible HG. 2004. Descending control of persistent pain: Inhibitory or facilitatory? Brain Research Reviews 46: 295–309. https://pubmed.ncbi.nlm.nih.gov/15571771/ (Accessed June 22, 2021).

Vianna DML, Landeira-Fernandez J, Brandão ML. 2001. Dorsolateral and ventral regions of the periaqueductal gray matter are involved in distinct types of fear. Neuroscience & Biobehavioral Reviews 25: 711–719. http://www.sciencedirect.com/science/article/pii/S0149763401000525 (Accessed June 14, 2015).

